# Proteomics and constraint-based modelling reveal enzyme kinetic properties of *Chlamydomonas reinhardtii* on a genome scale

**DOI:** 10.1101/2022.11.06.515318

**Authors:** Marius Arend, David Zimmer, Rudan Xu, Frederick Sommer, Timo Mühlhaus, Zoran Nikoloski

**Affiliations:** Bioinformatics, Institute of Biochemistry and Biology, University of Potsdam, Potsdam, Germany; Systems Biology and Mathematical Modelling, Max Planck Institute of Molecular Plant Physiology, Potsdam, Germany; Bioinformatics and Mathematical Modeling Department, Center of Plant Systems Biology and Biotechnology, 4000 Plovdiv, Bulgaria; Computational Systems Biology, TU Kaiserslautern, 67663 Kaiserslautern, Germany; Molecular Biotechnology & Systems Biology, TU Kaiserslautern, Kaiserslautern, Germany

**Keywords:** Turnover numbers, Metabolic modelling, Plant biology, Metabolic engineering

## Abstract

Biofuels produced from microalgae offer a promising solution for carbon neutral economy, and integration of turnover numbers into metabolic models can improve the design of metabolic engineering strategies towards achieving this aim. However, the coverage of enzyme turnover numbers for *Chlamydomonas reinhardtii*, a model eukaryotic microalga accessible to metabolic engineering, is 17-fold smaller compared to the heterotrophic model *Saccharomyces cerevisiae* often used as a cell factory. Here we generated protein abundance data from *Chlamydomonas reinhardtii* cells grown in various experiments, covering between 2337 and 3708 proteins, and employed these data with constraint-based metabolic modeling approaches to estimate *in vivo* maximum apparent turnover numbers for this model organism. The gathered data allowed us to estimate maximum apparent turnover numbers for 568 reactions, of which 46 correspond to transporters that are otherwise difficult to characterize. The resulting, largest-to-date catalogue of proxies for *in vivo* turnover numbers increased the coverage for *C. reinhardtii* by more than 10-fold. We showed that incorporation of these *in vivo* turnover numbers into a protein-constrained metabolic model of *C. reinhardtii* improves the accuracy of predicted enzyme usage in comparison to predictions resulting from the integration on *in vitro* turnover numbers. Together, the integration of proteomics and physiological data allowed us to extend our knowledge of previously uncharacterized enzymes in the *C. reinhardtii* genome and subsequently increase predictive performance for biotechnological applications.

**Significance statement:** Current metabolic modelling approaches rely on the usage of *in vitro* turnover numbers (*k_cat_*) that provide limited information on enzymes operating in their native environment. This knowledge gap can be closed by data-integrative approaches to estimate *in vivo k_cat_* values that can improve metabolic modelling and design of metabolic engineering strategies. In this work, we assembled a high-quality proteomics data set containing 27 samples of various culture conditions and strains of *Chlamydomonas reinhardtii*. We used this resource to create the largest data set of estimates for *in vivo* turnover numbers to date. Subsequently, we showed that metabolic models parameterized with these estimates provide better predictions of enzyme abundance than those obtained by using *in vitro* turnover numbers.

## Introduction

Microalgae can synthesize a wide range of high-value compounds (1) and biofuel precursors (2, 3) using industrial waste products and light energy, rendering them a key biotechnological resource propelling the transition to a net-zero carbon economy (4). However, economic feasibility of photosynthetic bioreactors requires further optimization of desired biotechnological objectives (4). Our ability to rationally engineer metabolism for biotechnological applications scales with our understanding of metabolism of organisms used as cell factories. Genome-scale metabolic models (GEMs), as mathematical representations of knowledge about metabolism, along with constraint-based modeling have facilitated the design of metabolic engineering strategies (5). Moreover, enzyme-related constraints, that rely on turnover numbers (*k_cat_*), have been shown to accurately predict various phenotypes including overflow metabolism (6–8), even without the usage of measurements of uptake fluxes (8). Further, these protein-constrained GEMs (pcGEMs) have been used to successfully identify engineering targets for biotechnological applications, such as an increased lysine production in *E. coli* (9).

The *k_cat_* data used in most pcGEM studies are obtained by laborious purification of the enzyme of interest and quantifying its maximum catalytic efficiency in an *in vitro* experiment (8). For organisms with available quantitative proteomic and physiological data, it is also possible to estimate the maximum apparent catalytic rate (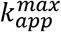) of an enzyme *in vivo* using constraint-based modelling (8). While it has been shown that *k_cat_* and 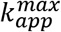 values are concordant for *E. coli* (10), for eukaryotic organisms like *S. cerevisiae* (11) and *A. thaliana* (12) lower correlation values between *k_cat_* and 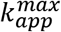 have been reported. This raises the question about the extent to which *in vitro* data can describe *in vivo* enzyme properties, particularly in eukaryotes. Nevertheless, most pcGEMs constructed to date rely on turnover numbers compiled in the public databases, such as: BRENDA (13) and SABIO-RK (14). While these databases offer comprehensive kinetic data for *E. coli*, only 10% of the entries in the union of the two databases cover enzymes of the Viridiplantae taxon. Further, the databases contain a total of only 85 turnover numbers (0.0012% of entries) specific to green algae. Thus, in order to make the powerful pcGEM modelling framework available to this biotechnologically relevant taxon, we must substantially increase the knowledge of *in vivo* turnover numbers.

Here we used cutting-edge mass spectrometry techniques (15, 16) to acquire a comprehensive set of protein abundance values from cultures of *C. reinhardtii* wild type and mutant strains grown under various conditions. We used this data set together with the recently developed minimization of non-idle enzyme (NIDLE) approach (17) to estimate 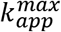 values for reactions catalyzed by single enzymes as well as decomposing the contribution isoenzymes to their catalyzed reactions, thus extending the state-of-the-art for estimation of 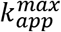 values by constraint-based modeling. Due to the sensitive proteomics approach we achieved a higher enzyme coverage than in previous works (10–12), extending the available literature data on *C. reinhardtii* by ~ 10-fold. In total, we obtained 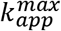 values for 568 including 46 transport reactions whose transport capacities are not quantifiable with current *in vitro* techniques. Our subsequent analysis corroborated the low correspondence between *k_cat_* and 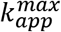 values in eukaryotic organisms. In line with these results, we showed that the substitution of *k_cat_* values in pcGEMs of *C. reinhardtii* with 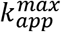 estimates improved predictive accuracy of enzyme resource allocation for unseen test conditions.

## Results and discussion

### High-quality protein abundance data from various experimental set-ups enable 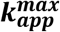 estimation

To obtain 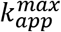 values for *C. reinhardtii* we employed comprehensive, high-quality proteomics data set encompassing 27 samples from various strains and growth conditions sampled at steady state. The absolute protein abundance data were generated based on the QConCATapproach (15, 16). QConCAT employs an isotopically labelled artificial protein containing concatenated peptides of multiple endogenous proteins as external standard to allow for absolute quantification of protein abundance. Using the concatamer to obtain a calibration curve we were able to obtain absolute protein quantification for up to 3708 (median: 3376) proteins (**Supp. Table 1**). On average, 28% of the measured proteins were annotated as enzymes and were included in the iCre1355 genome-scale metabolic model (GEM) of *C. reinhardtii* (**Fig. 1a**). In total, 936 of the 1460 proteins (64%) included as enzymes in iCre1355 were quantified in at least one experimental condition. The gathered data set covers about 100 enzymes more than the data used by Chen et al. (11); thus, this study marks the investigation of enzyme kinetic parameters from proteomics data with highest enzyme coverage to date.

**Fig. 1.**
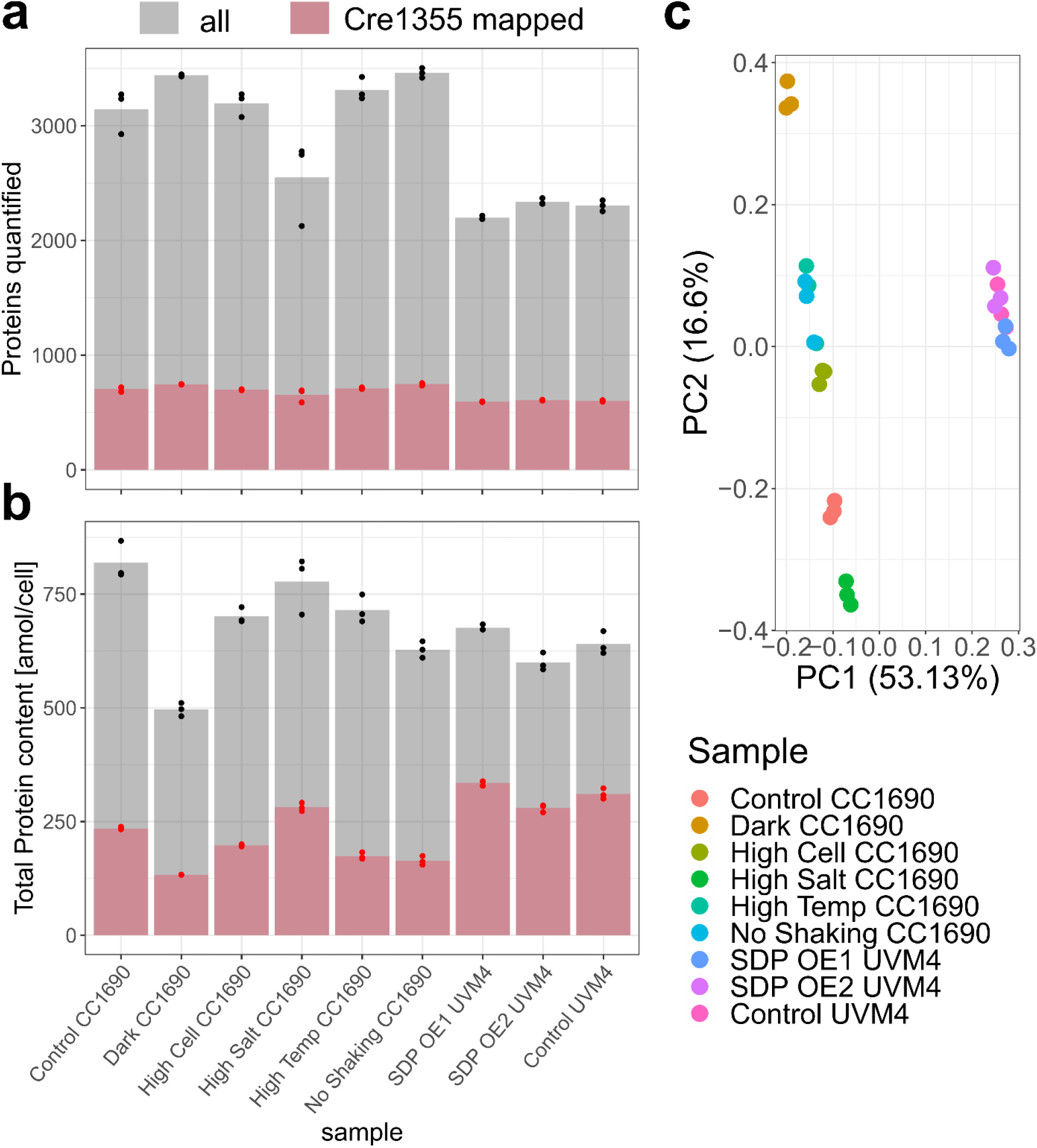
QConCATdata include various protein expression states and provide similar enzyme coverage. (**a**) Number of proteins quantified in at least two of the three replicates per condition, specified in the x-axis. (**b**) Principal component analysis of log-transformed abundance values of enzymatic proteins in *C. reinhardtii*. All replicates in the data set are plotted. (**c**) Total protein content summed over all proteins. In panels **b** and **c**, the dark red bar illustrates the number corresponding to enzymes present in the Cre1355 model.

We observed a smaller number of quantified proteins in the UVM4 strains compared to CC1690 (**Fig. 1a**). However, in terms of total quantified protein amount there is no systematic difference between the analyzed strains (**Fig. 1b**). Principal component (PC) analysis of the enzymatic proteins quantified in all samples reveals that replicates cluster together and experiments separate according to strain and culture conditions (**Fig. 1c**). The first PC resolves strain-specific effects and captures the majority of variance in the data set, while the second PC captures effects specific to the culture condition. Therefore, we concluded that the enzymatic proteins quantified here provide a wide and non-redundant set of *C. reinhardtii’s* metabolic states.

### Improved coverage of 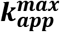 estimates for *C. reinhardtii*

Our main aim is to make use of the proteomics data to extend the sparse knowledge of enzyme kinetic properties in *C. reinhardtii*. To calculate apparent catalytic rates on a genome scale we used the NIDLE approach that minimizes the number of idle enzymes (i.e. those that do not carry flux, but have abundance measured), representing the principle of effective usage of cellular resources (17). NIDLE does not rely on maximizing growth as a cellular objective, but rather includes constrains from measured specific growth rates. It is formulated as a mixed-integer linear program (MILP) and does not enforce any proportionality between the measured enzyme abundance and reaction flux. The condition-specific flux distributions obtained by this MILP formulation are then used together with the absolute protein quantification to calculate the apparent catalytic rates, following established approaches (8, 17).

Here we expanded on the original NIDLE formulation to calculate estimates of isoenzyme *k_app_* values using a linear or quadratic formulation (see **Methods**). Based on this extension we were able to determine enzyme kinetic data for 18 and 41 reactions with multiple expressed isozymes based on the linear and quadratic formulations, respectively (**Supp. Fig. 1**). We decided to use the *k_app_* estimates of the quadratic formulation in the following analyses due to the higher coverage. In total, we obtained apparent catalytic rates for 568 enzyme catalyzed reactions in at least one of the experimental conditions (**Fig. 2a,b, Supp. Table 2**), which is the largest set of organism-specific *k_app_* estimates generated to date. The previously published pFBA (10) approach together with the QP for isoenzyme *k_app_* calculation only resulted in 489 estimates (**Supp. Fig. 2a**), that were highly correlated with the NIDLE results (Spearman correlation of log transformed values: 0.96, p<0.0001, n=483; **Supp. Fig. 2b**). Furthermore, in the NIDLE output for 52% of reactions we were able to calculate *k_app_* values in more than half of the investigated conditions. We observed that the largest group (n=189) of *k_app_* values was obtained from all nine considered conditions (**Supp. Fig. 3a**). These results gave us confidence that the maximum over the *k_app_* values for a reaction can serve as a good approximation of the *in vivo* turnover number. Upon determining 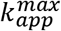, we observed the CC1690 and UVM4 standard mixotrophic growth conditions contributed the largest number of reactions operating at the maximum catalytic rate (**Supp. Fig. 3b**). Furthermore, there is no condition that does not contribute information to the calculated 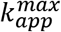 values. The distinction between samples based on their contribution to *k_app_* values is further supported by the principal component analysis using these values (**Supp. Fig. 3c**). In contrast to the principal component analysis based on the enzyme proteomic data, the largest difference between samples is observed between mixotrophic and heterotrophic growth conditions, resolved by both plotted principal components, while difference between mixotrophic samples is mainly explained by the second principal component (**Supp. Fig. 3c**).

**Fig. 2.**
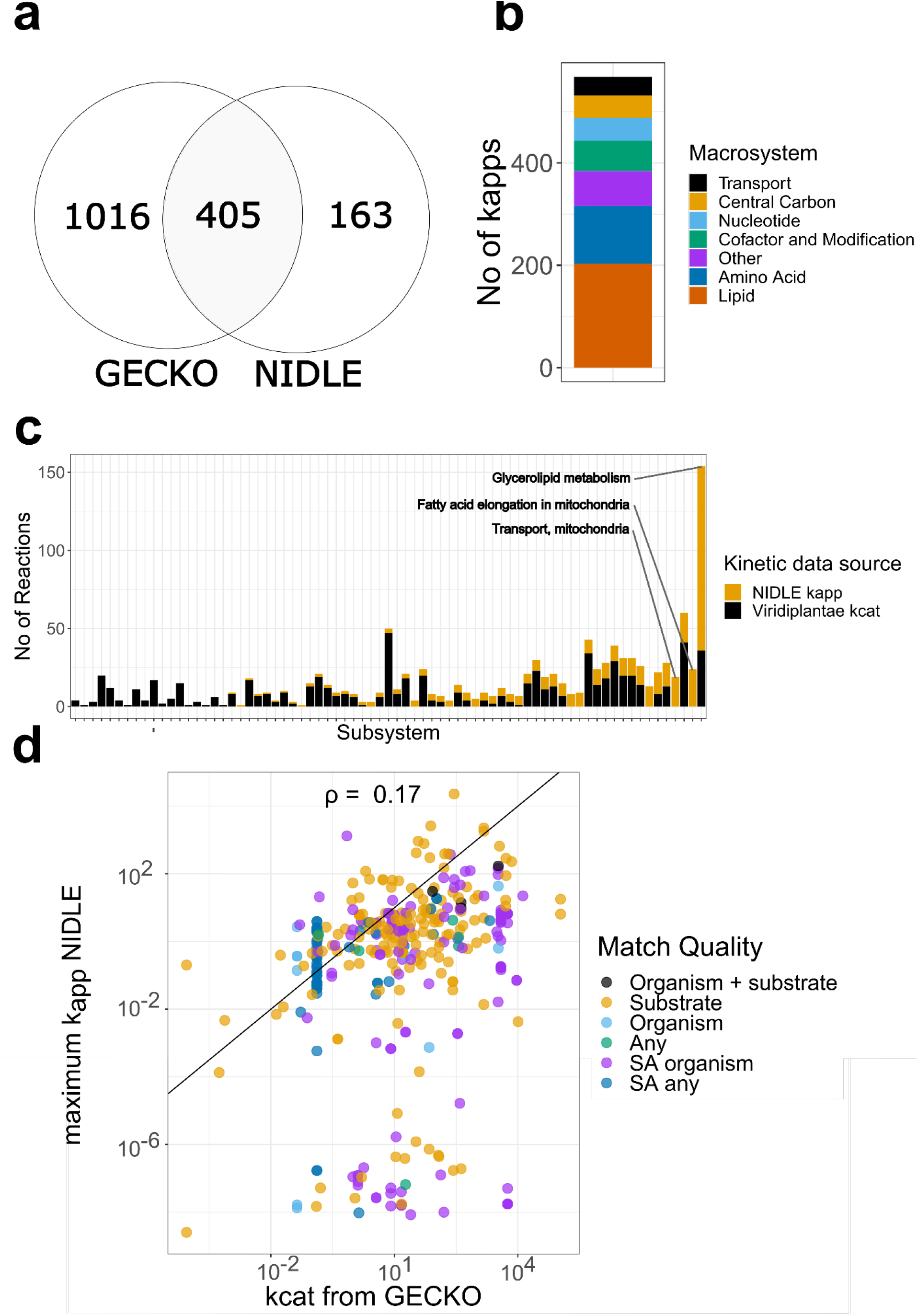
NIDLE combines physiological and proteomic data from experimental set-ups to obtain estimates of 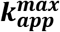. (**a**) Venn diagram showing the overlap in enzyme catalyzed reactions with maximum *k_app_* determined from NIDLE compared and *k_cat_* assigned based on EC Numbers by the GECKO heuristic (**b**) Stacked barplot indicating the number of 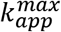 values that were determined in the different metabolic subsystem of iCre1355 GEM of *C. reinhardtii*. (**c**) The number of reactions with data on *k_cat_* from the Viridiplantae taxon have are indicated by a black bar, for each metabolic subsystem in iCre1355 (19). The stacked yellow bar indicates the extension of reactions for which 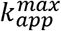 value was determined by NIDLE. (**d**) Scatterplot of the respective values in the intersection presented in panel a. The color code gives the matching criteria of *k_cat_* values from the GECKO heuristic in order of decreasing quality. SA: Specific activity.

The set of 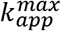 values presented here includes reactions from all major subsystems of primary metabolism (**Fig. 2b**), thus extending the current data on turnover numbers specific for *C. reinhardtii* available at BRENDA (13) and SABIORK (14) about ten-fold. Further, for 448 of the reactions with assigned 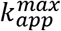 a query to these databases did not result in any known values in the whole Viridiplantae taxon. When we ranked the metabolic subsystems for which our data provide new enzyme kinetic information, we observed that the largest extension (for Viridiplantae-specific enzymes) was obtained for glycerolipid synthesis and mitochondrial fatty acid elongation (**Fig. 2c**). Aside from substantially increasing the kinetic information available for this photosynthetic organism, we also provide estimates of maximum catalytic rate for enzymes that are practically inaccessible to *in vitro* methods, because they are very difficult to purify and the measurement of reaction rate demands advanced assays. Namely, we were able to determine 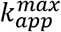 for 46 transport reactions (top subsystem “Transport, mitochondrial”, **Fig. 2c**) and their respective transporter proteins. Thus, our results provide valuable input for pcGEMs that currently cannot be obtained from existing databases.

### *k_cat_* values compiled by GECKO show no correspondence to the estimated 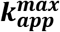

Studies in *S. cerevisiae* (11) and *A. thaliana* (12) found that *in vitro* determined turnover numbers provide a rather poor proxy of *in vivo* turnover numbers. Thus, we were interested to identify if curated literature *k_cat_* values for *C. reinhardtii* correspond to the determined *in vivo* 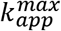 values. We used the recently updated GECKO heuristic (7, 18) to assign the phylogenetically closest available *k_cat_* values from BRENDA (13) to reactions (**Fig. 2a**). For the overlap of reactions that where assigned a maximum catalytic rate by both GECKO and NIDLE, we found that our results corroborate the findings from the two eukaryotes. More specifically, the correspondence between log-transformed values is low (Spearman correlation of 0.19, p-value < 0.001, n=405). Moreover, the *in vivo* 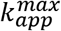 values are systematically lower than the corresponding literature turnover number (**Fig. 2d**). Since the aim of the GECKO approach is to parameterize as many reactions as possible, it iteratively relaxes the matching criteria when assigning *k_cat_* values from literature. While we observe a higher correspondence for *k_cat_* values from endogenous *C. reinhardtii* proteins (Spearman correlation 0.75, p-value=0.0019, n=14), there is no obvious difference in the other groups (**Fig. 2d**). The scatterplot also reveals that GECKO assigns many reactions in the lowest quality group the same *k_cat_* value, while NIDLE provides specific 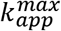 values ranging several orders of magnitude (**Fig. 2d**). These results underline once more that literature turnover numbers are a suboptimal source of parameters for pcGEMs, due to the *in vivo/in vitro* effects and problematic matching of organism unspecific kinetic data.

### Parameterization of pcGEMs with the estimates of *in vivo* 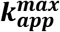 values show improved enzyme usage prediction

To investigate if the 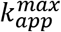 values calculated from NIDLE result in an improvement of the predictive performance of pcGEMs we generated a mixotrophic and a heterotrophic pcGEM for *C. reinhardtii* based on the models published by Imam et al. (19) using the GECKO toolbox (18). In a first step we used the chemostat data set of Imam et al. to test the effect of the obtained 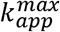 values on growth rate predictions. For each tested condition a so-called raw GECKO model was built, including the *k_cat_* values extracted from literature. The over-constraining *k_cat_* values were then corrected using the objective control coefficient heuristic and the average enzyme saturation coefficient, *σ*, was fitted according to the measured growth rate (19) (**Supp. Table 3**). To obtain pcGEMs using the NIDLE 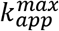, the *k_cat_* values in both the raw and the corrected GECKO models were substituted with the respective 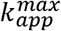 estimates, were available. When we compared pcGEM model predictions with flux balance analysis (FBA) and experimental measured growth rate, we found that raw GECKO models with and without usage of 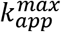 underestimate growth compared to FBA predictions (**Fig. 3a**). The only exception was the heterotrophic conditions, in which NIDLE raw pcGEM predicts higher growth then experimentally observed. In all cases, the experimental growth rate was reached only after the *k_cat_* correction step and refitting *σ*. For heterotrophic conditions, *σ* was fitted to ~ 0.4 of the value in autotrophic in mixotrophic conditions (**Supp. Table 3**), indicating that in heterotrophic growth many enzymes are expressed considerably higher than necessary to maintain metabolic flux (**Fig. 3a**). Comparing the performance of NIDLE pcGEMs we did not observe a strong effect on growth rate predictions. As expected, the prediction error was higher than in the corrected GECKO pcGEMs, since the latter were fitted to the experimental growth rate; however, the introduced error (RMSE 0.0163) was comparable to that of canonical FBA (RMSE 0.0183) (**Fig. 3a**).

**Fig. 3.**
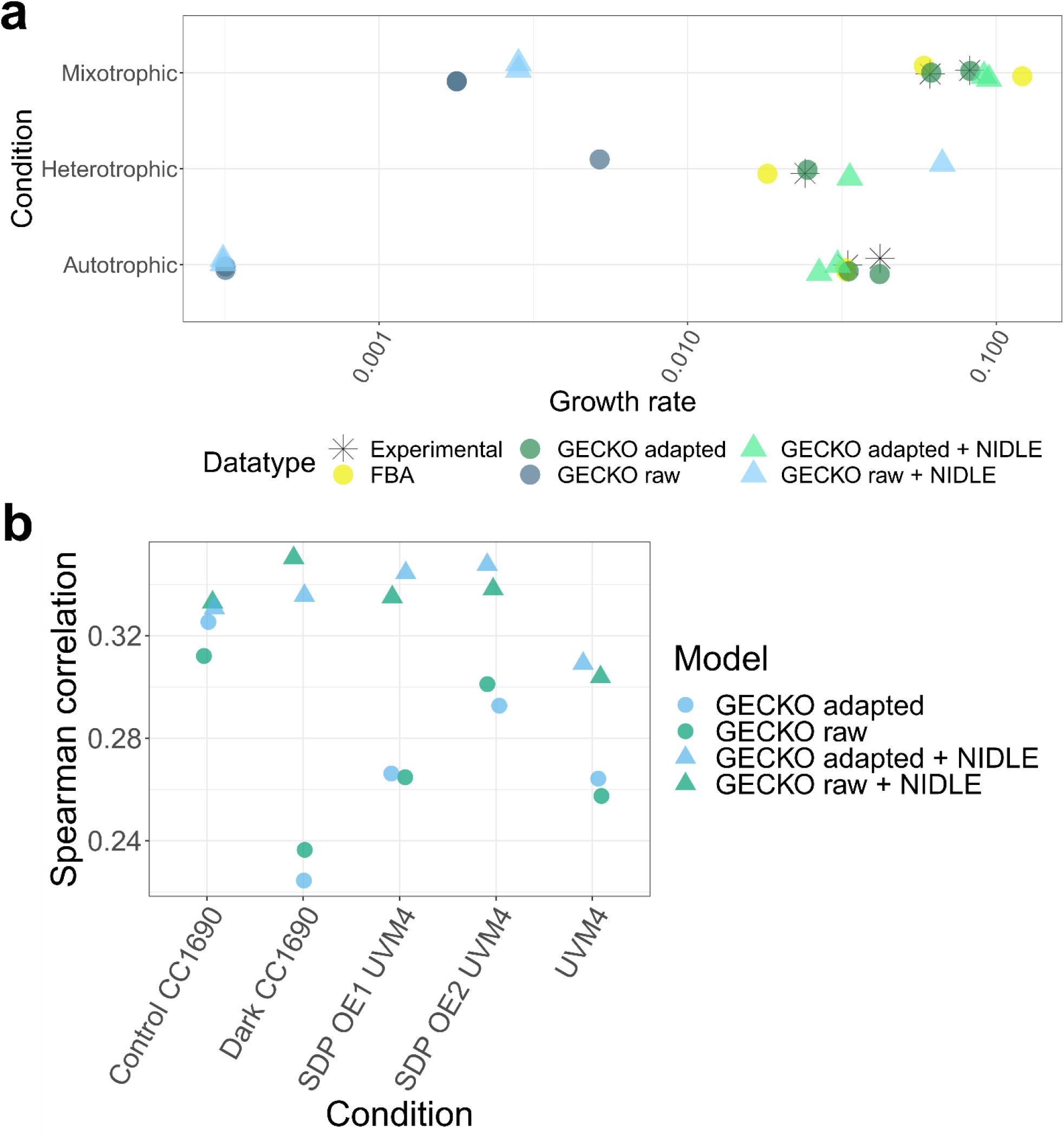
The usage of 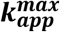 increases the prediction accuracy of enzyme usage for unseen experiments. (**a**) Comparison of experimental data from chemostat cultures (19) and predictions from FBA and pcGEMs parameterized with uncorrected *k_cat_* values obtained from BRENDA (13) (GECKO raw), corrected *k_cat_* values using GECKO heuristic (GECKO adapted) or updated with enzyme wise 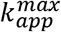 from NIDLE (+ NIDLE) (**b**) Spearman correlation of predicted enzyme usage based on pcGEMs and observed enzyme abundance in QConCat data set. The tested condition was not considered when calculating the 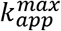 values from NIDLE.

Since the maximum catalytic capacity used in pcGEMs quantifies the enzymatic expenditure to support a certain reaction flux, another important application of these models is in predicting the allocation of total enzyme mass into specific enzymes. Therefore, we tested the effect of the 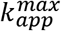 values obtained by NIDLE on the accuracy of enzyme usage prediction in the standard conditions included in our proteomics data set. To allow for an informative comparison, we left out the NIDLE *k_app_* values calculated in the tested condition when calculating 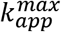 for the respective NIDLE pcGEMs (see **Methods**). We predicted enzyme usage coefficients by fixing the flux through biomass reaction to the experimentally observed growth rate and minimizing the total enzyme mass. Next, we calculated the Spearman correlation between these predictions and the measured enzyme abundances. Interestingly, we observed that models containing 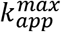 from independent proteomics samples showed higher predictive performance in the unseen condition than the canonical GECKO models (**Fig 3b**). This observation was made irrespective of the tested condition or whether the raw or corrected GECKO model was compared, even though the corrected GECKO model is fitted to the experimental growth rate from mixotrophic chemostat experiments. Thus, we were able to demonstrate that the NIDLE approach successfully uses information from physiological data to calculate maximum catalytic capacity values which are a better predictor of enzyme usage than widely used literature values.

## Conclusion

We presented a protein abundance data set with extensive coverage of the proteome response to various perturbations. This data set comprises a valuable resource for systems biology studies in *C. reinhardtii*. Here we made use of this resource to considerably expand the information for green algae. Due to the extended NIDLE formulation we were able to estimate 568 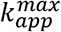 values for enzyme catalyzed reactions, compared to 436 and 358 previously reported 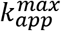 values in *E. coli (10)* and *S. cerevisiae* (11), respectively. The obtained information allows to quantify the costs for different cellular pathways and thus fosters the application of advanced metabolic engineering strategies in the biotechnologically relevant taxon of green algae.

## Materials and methods

### Data set assembly

Analyzed *Chlamydomonas reinhardtii* (*C. reinhardtii*) data sets included data from previously published QConCAT studies available under PRIDE (20) data set identifier PXD018833 (Control UVM4, SDP OE1 UVM4, SDP OE2 UVM4) and were further augmented by data sets measured as part of this study and made publicly available under the PRIDE identifier PXD037599 (Control CC1690, Dark CC1690, High Cell CC1690, High Salt CC1690, High Temp CC1690, No Shaking CC1690). As control cultivation conditions for *C. reinhardtii* CC1690 cells, cultures were grown for 48 hours in Tris-Acetate-Phosphate (TAP) medium using a rotatory shaker operating at 2 turns per second, while being constantly illuminated at 100 μmol photons m-2 s −1 at and held at 24°C. Data sets differing from control conditions were created by alteration of listed growth parameters (a detailed description of modified factors is available in **Supp. Table 4**).

### LC-MS/MS measurement and raw data analysis

After cell harvesting and protein extraction, all samples were spiked with a master mix of Chlamydomonas-specific QConCAT proteins, digested tryptically, and analyzed via LC-MS/MS (Eksigent nanoLC 425 coupled to a TripleTOF 6600, ABSciex) as described in Hammel *et al.*, 2020 (16). Quantitative analysis of MS/MS measurements was performed using ProteomIQon 0.0.7 (21). Peptide searches were performed upon the assembly of a peptide database based of the *Chlamydomonas* proteome based on version JGI5.5 of the *C. reinhardtii* genome blended with the sequences of spiked-in QconCAT proteins. The search space included methionine oxidation and acetylation of protein N termini as variable modifications and was extended by ^15^N-labeled variants of *Chlamydomonas* proteins. False discovery rate thresholds for peptide spectrum matches and protein group identifications were set to 1%. Following peptide spectrum matching, ion abundances were estimated by integration of the XIC area.

### QConCAT-based estimation of absolute protein abundances

To obtain absolute protein abundances, we first aggregated ion species to the modified peptide level by summation (e.g., different charge states). Differently modified versions of the peptides were aggregated to the peptide and then protein-group level by median-based aggregation, yielding preliminary protein abundance estimates. Computing the ratio between native, unlabeled peptides and ^15^N-labeled peptides originating from spiked-in QConCAT proteins allowed to estimate absolute protein abundances for a multitude of different *C. reinhardtii* proteins (16), as previously described. With these high-quality QconCAT-based abundance measurements at hand, we were able to create calibration curves by regressing the latter on the preliminary protein abundance estimators, and thus to compute proteome-wide absolute abundance estimates.

### Processing of non-proteotypic peptides

A data set entry is considered to have ambiguous entries if its quantification is based on a peptide that is non proteotypic (i.e. is present in multiple proteins); otherwise, the protein is defined to have an unambiguous entry. For ambiguous entries, an iterative approach was used to remove them in each sample. If an unambiguous entry of one of the mapped proteins was present, the corresponding concentration was subtracted from all ambiguous entries of this protein and the protein ID was removed from the ambiguous entries. If the cellular concentration of an ambiguous entry was smaller than 0 after subtracting, the entry was removed from the sample. This procedure was repeated until no further protein IDs could be removed from ambiguous entries. The remaining ambiguous entries were removed from the sample data. Proteins that were only quantified in one of the three biological replicates were removed from the data set. For the remaining data the median over the measured replicates was used in the downstream analyses.

### GEM used in constraint-based modeling

The most recent SBML and COBRA compatible model of *Chlamydomonas reinhardtii* “iCre1355” (19) was employed. The erroneous reaction formular of ‘CAT’ was updated to “ 2 h2o2[c] -> 2 h2o[c] + o2[c]”. GPR rule syntax was updated to not include “… and (GENE1 or GENE2 …) …” rules. All model modifications and mathematical programs solved in this study was carried out using the COBRA toolbox (22) and GUROBI solver (23) in MATLAB (24).

### NIDLE

We used the iCre1355 mixotrophic and heterotrophic model with the respective culture conditions used in the proteomics experiments. The NIDLE approach is formulated for positive, real valued flux variables; therefore, the models were converted to irreversible by splitting each reversible reaction into irreversible forward and backward reactions. All uptake reactions were constrained by the model supplied bounds except for acetate uptake. For mixotrophic conditions a linear regression model was fitted based on the mixotrophic chemostat culture measurements from Imam et al. (19), in which acetate uptake rate was predicted based on growth rate. The model was fitted using the R *lm* function(25) with default options, and the model predicted uptake rate increased by the standard error of prediction was used as an upper bound on the acetate uptake rate for the mixotrophic culture scenarios (19). For the heterotrophic condition the maximum measured acetate uptake rate from the Imam et al. heterotrophic chemostat data was set as an upper bound (**Supp. Table 5**). We adapted the source code in the NIDLE repository (17) to the iCre1355 model, but the formulation of the NIDLE approach, based on a mixed-integer linear program, remained unchanged.

To calculate *k_app_* values for homomeric isoenzyme catalyzed reactions we first determined if only one of the catalyzing isoenzymes is quantified in the respective condition. If this was the case the *k_app_* was calculated in the same way as for the homomeric enzymes, i.e. the reaction flux of reaction *i* in condition *j* divided by the respective enzyme abundance *E* gives the apparent catalytic rate,

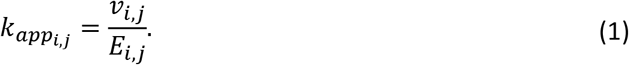

To convert enzyme abundances measured in amol/cell to mmol/gDW the literature cell dry weight of 48.000 pg (26) was used for a mixotrophic grown cell and the dry weight of other conditions was calculated from measured cell volume assuming constant dry weight density.

In the case that multiple isoenzymes have been measured by mass spectrometry we integrated information from different conditions to decompose the contribution of different isoenzymes to the observed flux. We took advantage of the fact that in the mixotrophic standard growth conditions best approximate the maximum apparent catalytic rate for the majority of enzymes (**Supp. Fig. 2c**), and assumed equal *k_app_* values for an isoenzyme in the different conditions. This allowed us to formulate a quadratic problem based on the flux predictions and enzyme abundance measurements in the four mixotrophic standard conditions (i.e. Control CC1690, Control UVM4, SDPOE1 UVM4, SDPOE1 UVM4, SDPOE2 UVM4), given in the following

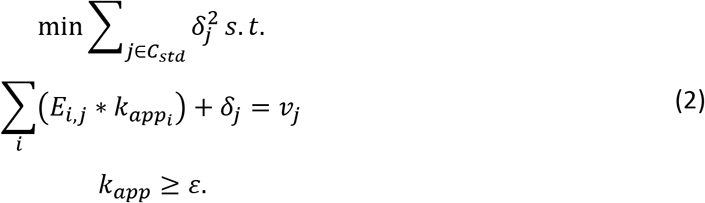

More specifically, we obtain *k_app_* estimates by minimizing the quadratic sum of residuals between flux supported by the *k_apps_* and obtained from NIDLE, over all conditions j. We chose *ε* of 10^−10^ · 3600 [h^−1^] since both the smallest turnover number in the joint public databases (5.8 * 10^−10^[s^−1^]) (13, 14) and calculated from homomeric reactions (4.0 * 10^−10^[s^−1^]) were in this order of magnitude. The effective reaction-specific *k_app_* for each condition was then calculated as the average weighted by the protein abundance in the given condition,

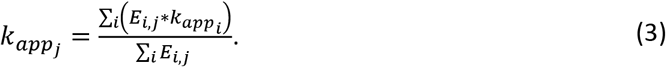

We also compared the solution from minimizing the ℓ_1_-norm of the error term, *δ*,

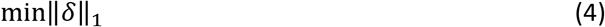

subjected to the same constrains (**Supp. Fig. 1a**). We did not consider *k_app_* values equal to the lower bound, *∊*.

### pcGEM creation using the GECKO toolbox

The GECKO toolbox (7, 18) was used to integrate maximum catalytic rate data into a pcGEM. Based on each of the published models (mixo-auto-, and heteroptrophic), and the obtained chemostat experiments (19) corresponding pcGEMs were created. Scripts were adapted according to the README instructions (https://github.com/SysBioChalmers/GECKO). For compatibility with the GECKO toolbox, the JGI gene ids in iCre1355 were converted to Uniprot ids and introduced duplicates where removed. The protein content used for biomass rescaling and limiting of the enzyme pool reaction was taken from the measurements of Boyle & Morgan (27). For all pcGEM simulations the uptake rate bounds of the macronutrients ammonium, phosphate, and carbon dioxide where set to 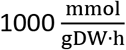. The average protein abundance over all sampled conditions was used to calculate the factor *f* (only proteins without missing values were used). Growth associated maintenance was not refitted. A corrected model based on the observed chemostat growth measurements in the model publication (19) was created using GECKOs objective control coefficient heuristic to correct over constraining *k_cat_* values, and the sigma factor was fitted. The NIDLE pcGEMs were generated by substituting GECKO-assigned *k_cat_* values for each enzyme pseudometabolite in the augmented stochiometric matrix with the maximum of all NIDLE obtained 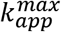 calculate over all reactions this enzyme catalyzes **(Supp. Table 6**). For the comparison of growth rate predictions the respective biomass reaction was used as objective function.

The pcGEM fitted with chemostat data from “Mixotrophic_Rep3” and “Heterotrophic_Rep1” were used for the simulation of enzyme usage in the proteomics experiments of the respective growth scheme. In the NIDLE models used for enzyme usage comparison the substituted 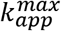 values were calculated omitting the *k_app_* values obtained from the tested condition. The same condition specific uptake flux constraints as in the NIDLE problems were used. Flux trough the biomass reaction was fixed to 0.99 of the observed growth rates and the flux through the “draw enzyme pool” reaction was minimized.

### Querying the BRENDA and SABIO-RK database

Turnover numbers of non-mutated enzymes together with organism and EC-number information were downloaded as text files from BRENDA (13) and SABIO-RK (14) databases (status 07/2022) and joint. For the reactions with EC-number annotation in iCre1355 (19) the following matching criteria for enzymes with fully matching EC-number where tried in the given order:

1. Chlamydomonas taxon & substrate
2. Chlamydomonas taxon
3. Viridiplantae taxon & substrate
4. Viridiplantae taxon

The maximum of all *k_cat_* values in the first criteria with non-zero number of matches was assigned as comparison *k_cat_*.

## Code and data availability

Code for the updated NIDLE approach and generation of the presented results is publicly available as GitHub repository:https://github.com/arendma/Crekapp.git. The proteomics data used for this study was deposited at PRIDE database (20) with identifiers PXD018833 (UVM4 data set) and PXD037599 (CC1690 data set).

