## Supplementary Figures for "Proteomics and constraint-based modelling reveal enzyme kinetic properties of *Chlamydomonas reinhardtii* on a genome scale"

**This PDF file includes:**

Supplementary Figures 1 to 3

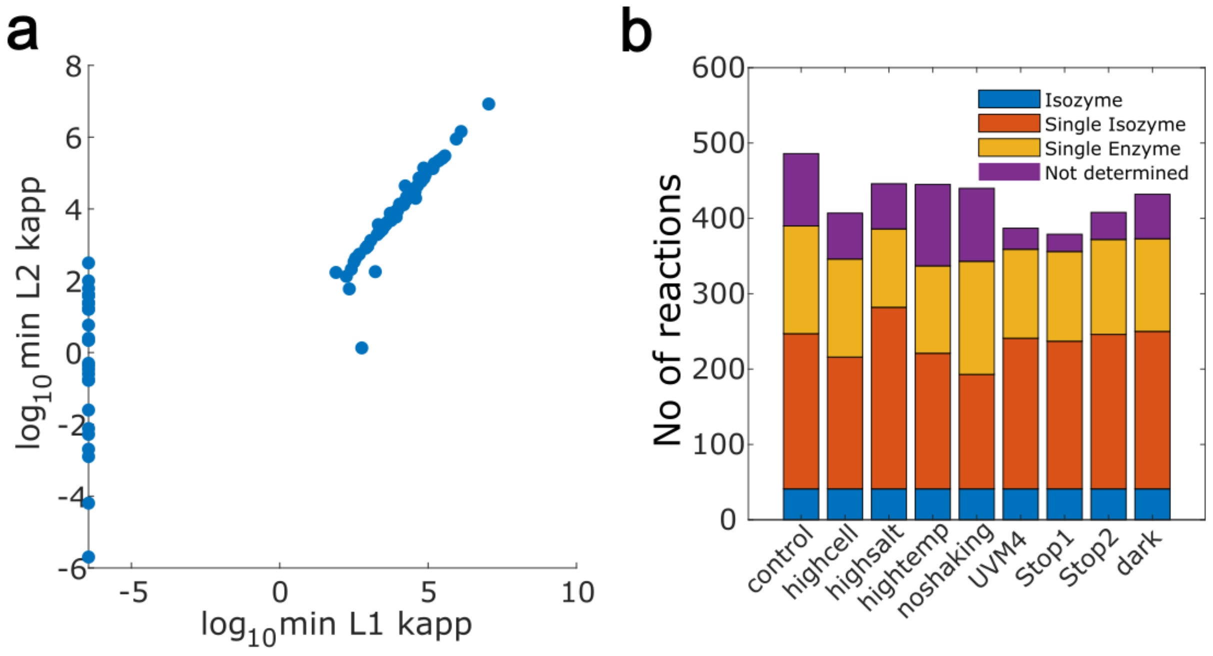


**Supplementary Figure 1. Properties of $k_{app}$ calculated for isoenzyme reactions. (a)** Scatterplot of enzyme-specific $k_{app}$ calculated for reactions with multiple expressed isoenzymes. The x-axis provides the estimates obtained by the linear formulation (i.e. minimizing the $\ell^1$- norm of the error term $\delta$). The minimal values correspond to the lower bound of $\epsilon = 3.6 \cdot 10^{-7}\ h^{-1}$. The y-axis provides the values estimated by the quadratic formulation. **(b)** Stacked barplot indicating the number of reactions for which single enzymes or (multiple) isoenzymes have been quantified. The category "not determined" originates from reactions with available protein abundance that do not carry flux.

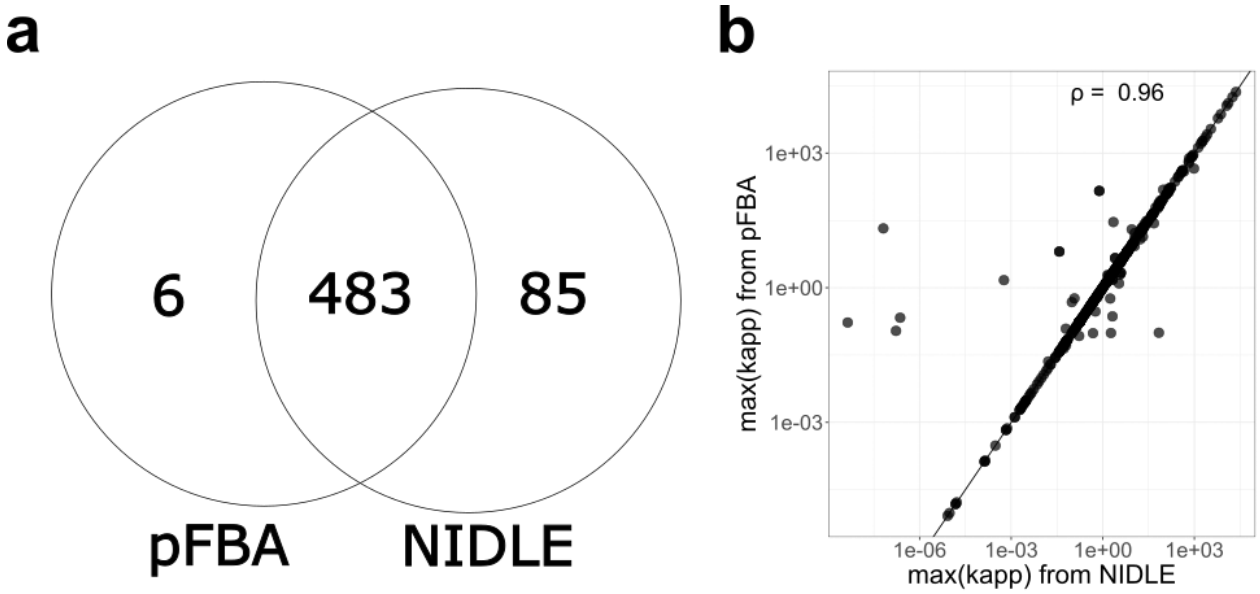


**Supplementary Figure 2. Comparison between pFBA and NIDLE based $\boldsymbol{k_{app}^{max}}$ estimates: (a)** Venn Diagram of enzyme catalyzed reactions for which $k_{app}^{max}$ could be estimated by the two approaches. **(b)** Scatterplot of $k_{app}^{max}$ values for the intersect in reactions. Values are plotted in log-scale. $\rho$: Spearman correlation of log-transformed values.

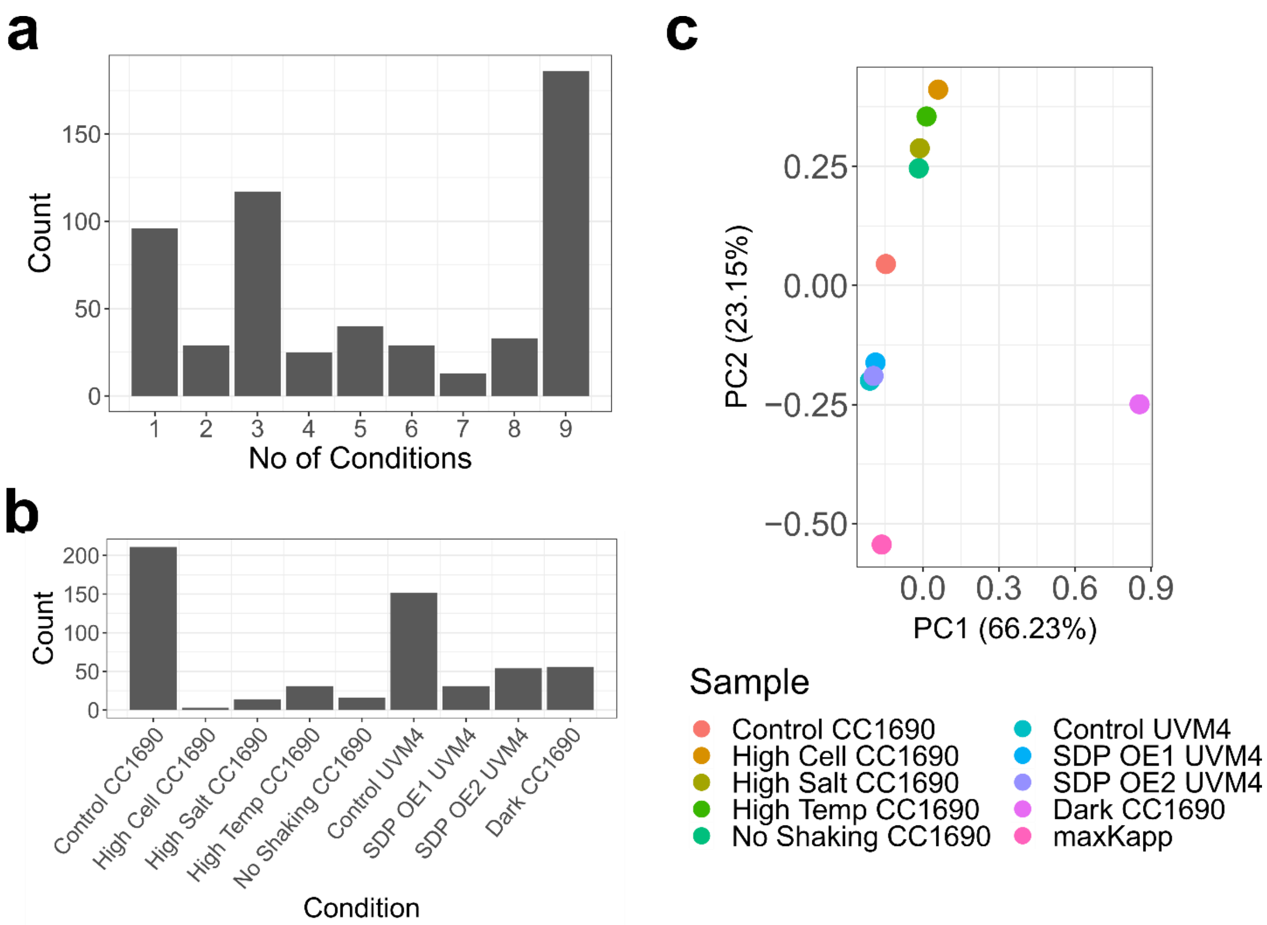


**Supplementary Figure 3. Distribution of $k_{app}$ values.** (**a**) Histogram of the number of conditions in which $k_{app}$ was calculated for the homomeric or isoenzyme enzyme catalyzed reactions in the iCre1355 model. (**b**) Number of homomeric or isoenzyme-catalyzed reactions for which the maximum observed $k_{app}$ was found in the respective condition. (**c**) Principal component analysis of log-transformed $k_{app}$ values per condition and the vector of maximum $k_{app}$ values. Only reactions that had non-zero values in all conditions were taken into account. Values were log-transformed before calculation of principal components.
